# Bioluminescence Assay of Lysine Deacylase Sirtuin Activity

**DOI:** 10.1101/2023.08.10.552871

**Authors:** Alexandria N. Van Scoyk, Orlando Antelope, Anca Franzini, Donald E. Ayer, Randall T. Peterson, Anthony D. Pomicter, Shawn C. Owen, Michael W. Deininger

## Abstract

Lysine acylation can direct protein function, localization, and interactions. Sirtuins deacylate lysine towards maintaining cellular homeostasis, and their aberrant expression contributes to the pathogenesis of multiple pathological conditions, including cancer. Measuring sirtuins’ activity is essential to exploring their potential as therapeutic targets, but accurate quantification is challenging. We developed ‘SIRT*ify*’, a high-sensitivity assay for measuring sirtuin activity *in vitro* and *in vivo*. SIRT*ify* is based on a split-version of the NanoLuc® luciferase consisting of a truncated, catalytically inactive N-terminal moiety (LgBiT) that complements with a high-affinity C-terminal peptide (p86) to form active luciferase. Acylation of two lysines within p86 disrupts binding to LgBiT and abates luminescence. Deacylation by sirtuins reestablishes p86 and restores binding, generating a luminescence signal proportional to sirtuin activity. Measurements accurately reflect reported sirtuin specificity for lysine acylations and confirm the effects of sirtuin modulators. SIRT*ify* effectively quantifies lysine deacylation dynamics and may be adaptable to monitoring additional post-translational modifications.

## INTRODUCTION

Reversible acylation of protein lysines regulates cellular processes, including transcription, metabolism, and signaling, to influence critical cell fate decisions, such as differentiation and apoptosis. To date, at least twenty different lysine acyl modifications have been reported, including acetylation, crotonylation, succinylation, and glutarylation^1^. Although recent data indicate that, with the exception of acetylation, non-enzymatic reactions account for the bulk of lysine acylations^1–5^, deacylation depends mainly on the activity of a single family of enzymes, termed sirtuins^2,3^.

The sirtuin family consists of seven members (SIRT1-7) with diverse and partially overlapping substrate specificity, cellular location, and function. Sirtuins are evolutionarily conserved NAD^+^-dependent lysine deacylases implicated in several fundamental biological processes^6,7^. Dysregulation of sirtuins is associated with a range of pathological conditions, including diabetes, inflammatory disorders, neurodegenerative diseases, and cancer. Considering their impact on key cellular processes, it is imperative to develop effective tools to characterize and monitor acyl modifications by sirtuins *in vitro* and *in vivo* to delineate their potential as therapy targets^6,8–13^.

Several assays have been developed to measure sirtuin activity, but only a few are commercially available. The most widely used test is FLUOR DE LYS® (FDL), a two-step assay that probes deacylation of an acyl-lysine peptide conjugated to aminomethylcoumarin (AMC) and a fluorescence quencher^14^. Following deacylation, trypsin is added as a ‘developing agent’ to cleave the AMC from the deacylated peptide-quencher, and the resulting increase in fluorescence is proportional to sirtuin activity. The utility of the FDL assay is limited by its tendency to produce false positive results from nonspecific interactions of the fluorophore with the target probe^15–19^ and incompatibility with more complex systems, including cell lysates, intact cells, and living organisms. Various other sirtuin assays have been reported that use chemical fluorescent and molecular self-assembly probes, radioisotope-labeled histones, nicotinamide release, or mass spectrometry^20–22^. These assays are neither widely used nor readily available, reflecting a range of limitations related to chemical stability, selectivity, expense, and/or toxicity^15–19,23–25^.

To address the limitations of current sirtuin assays, we developed SIRT*ify*, a luminescent assay for measuring and imaging lysine-deacylase activity in intact cells and *in vivo*. SIRT*ify* is based on a split version of NanoLuc^®^, an engineered luciferase from the deep-sea shrimp *Oplophorus gracilirostris*^26^. In the split-NanoLuc complementation system, removal of the C-terminal β-strand of the β-barrel domain renders NanoLuc catalytically inactive^26^. Acylation of the two lysines present within the small β-strand peptide fragment of split-NanoLuc reduces the complementation-based activity of the split-NanoLuc to very low levels, while deacylation by sirtuins restores luciferase activity. We demonstrate that SIRT*ify* accurately measures sirtuin activity in cell-free systems, *in vitro and in vivo*, and is useful for identifying sirtuin activity modulators without many of the issues seen in these complex systems with other assay formats.

## RESULTS

### Biochemical Characterization of Acylated Peptides

NanoLuc® is an engineered luciferase consisting of 11 antiparallel strands forming a β-barrel that is capped with four α-helices. The first split-NanoLuc complementation system demonstrated that removal of the C-terminal β-strand of the 10-stranded β-barrel renders NanoLuc catalytically inactive^26^. Sequence optimization of the C-terminal peptide identified a series of 11 amino acids peptides spanning five orders of magnitude in affinity for the large, truncated fragment of NanoLuc (LgBiT)^26^. From this series, we selected peptide ‘86’ (p86) because of its high affinity to LgBiT (K_D_=0.7×10^-9^ M) and because it contains two lysines located at position eight and nine of the peptide (NH_2_-VSGWRLF**KK**IS-OH)^26^. We hypothesized that ε-acylation of these lysines would reduce affinity to LgBiT, preventing reconstitution of active NanoLuc (Figure 1A, B). Further, removal of the respective acylation by a sirtuin should restore native p86, allowing for the reconstitution of active luciferase and corresponding luminescence.

**Figure 1.**
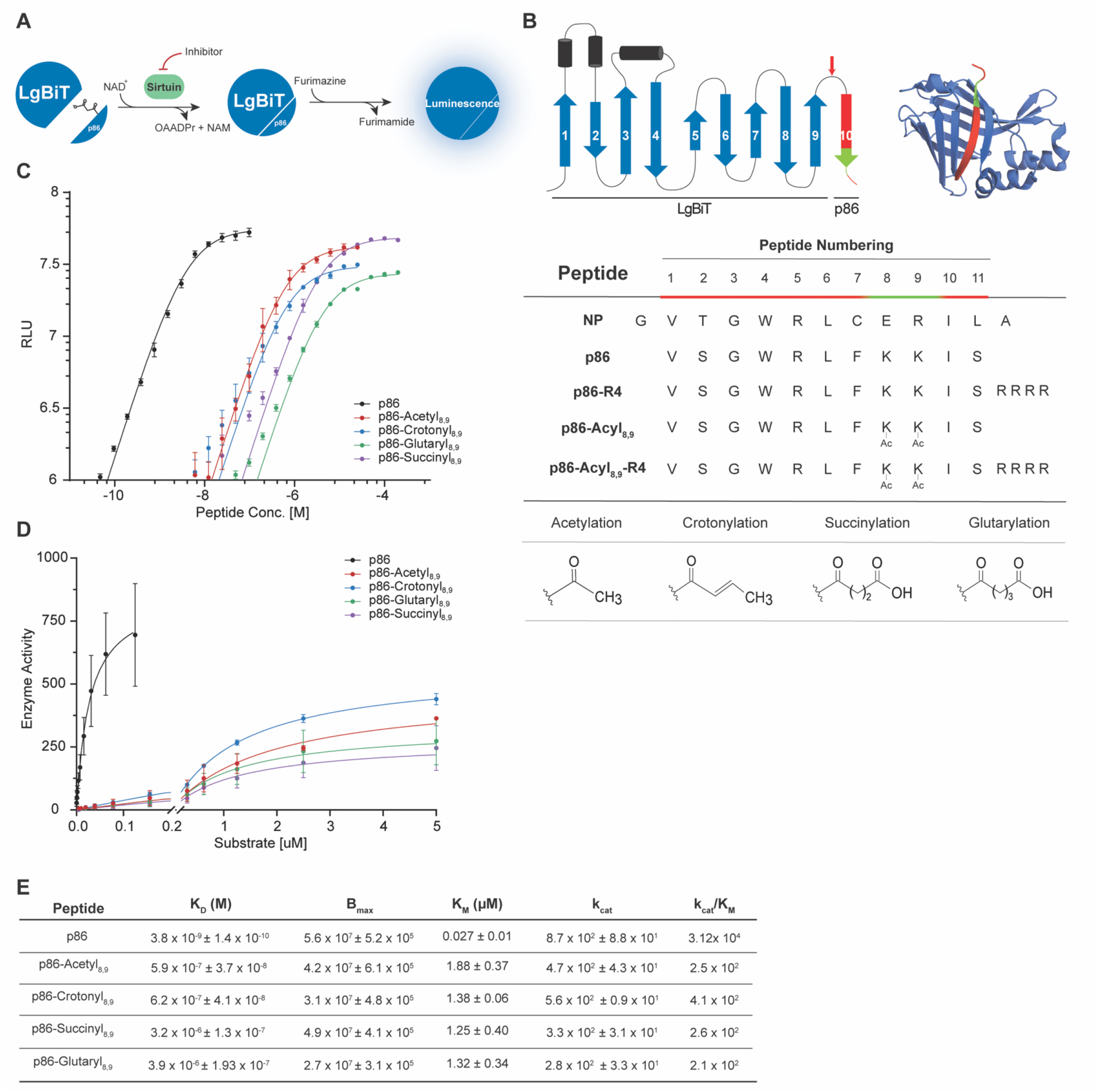
Design strategy of the luminescence-based lysine deacylase assay. (**A**) Schematic representation of the split-NanoLuc modified system adapted for detection of sirtuin activity by acylation of the lysine residues with peptide p86. NAD^+^ -nicotinamide adenine dinucleotide; NAM – nicotinamide; OAADPr – O-Acetyl-ADP-Ribose. (**B**) Structural representation of split-NanoLuc luciferase system, consisting of LgBiT (β-barrels 1-9) and p86 11-aa peptide (β-barrel 10). Sequences of 13-aa native peptide (NP), and p86 peptides (**C**) Titration of LgBiT with p86 and lysine-acylated p86 peptides. K_D_ values were estimated by fitting data to one site-specific binding model on GraphPad. (**D**) Kinetic parameters of furimazine with acylated peptides were estimated by fitting data to the Michaelis-Menten equation (**E**) Table of kinetic parameters shown in (**C**) and (**D**). Results in **C**–**E** are from independent experiments performed three times. Data show means with standard deviation.

As a proof of concept, we compared activity for unmodified, single, or dually acetylated and succinylated p86. Single acetylation or succinylation at either K8 or K9 resulted in only a minor decrease of 1.1 and 1.7-fold in luminescent signal, respectively (Supplemental Figure 1). In contrast, dual acetylation or succinylation at K8 and K9 resulted in a 5.8 and 7.5-fold decrease in signal. As the dual lysine modification was more effective in decreasing luminescence, we modified p86 peptides at both K8 and K9 for a series of acyl modifications--acetyl, crotonyl, succinyl, and glutaryl—to produce p86-Acetyl_8,9_, p86-Crotonyl_8,9_, p86-Succinyl_8,9_, and p86-Glutaryl_8,9_, respectively, which we collectively termed p86-Acyl_8,9_ peptides.

To determine changes in binding affinity induced by acylation, we measured the dissociation constants (*K_D_*) of LgBiT and p86-Acyl_8,9_ interactions by titration. We first determined the *K_D_* of p86 by fitting titration data to a standard Michaelis-Menten model and found it comparable to published data (3.80×10^-9^ M vs. 0.7×10^-9^ M reported by Dixon et al.^26^). We next determined *K_D_* values for p86-Acyl_8,9_ peptides (Figure 1C, E). Lysine acylation consistently decreased binding affinity for the large split-NanoLuc fragment of all p86-Acyl_8,9_ peptides. Compared to p86, p86-Acyl_8,9_peptides showed between ∼175-fold (p86-Acetyl_8,9_) and ∼1000-fold (p86-Glutaryl_8,9_) reduced affinity to LgBiT. Next, we measured kinetic parameters, including turnover number (*kcat)* and Michaelis constant (*K_M_*) of the p86-Acyl_8,9_ peptides relative to native p86. Under our assay conditions, *K_M_* of p86 with LgBiT was more than 60-fold lower than that of p86-Acyl_8,9_peptides (0.03 μM vs 1.25 - 1.88 μM, respectively). Accordingly, the catalytic efficiency of LgBiT expressed as (*kcat/K_M_*) decreased from 3.12×10^4^ μM^-1^s^-1^ for p86 to 2.10-4.08×10^2^ μM^-1^s^-1^ for p86-Acyl_8,9_ peptides (Figure 1D, E). Our results demonstrate that, although acylated peptides can still bind to LgBiT and enable substrate conversion, this activity is significantly reduced compared to native p86 peptide.

We tested whether sirtuin activity and specificity toward the p86-Acyl_8,9_ peptide substrates were maintained and whether luciferase activity was restored in the presence of sirtuins. Of the SIRT1-7 enzymes, we chose to focus on SIRT1, 2, 3, and 5 due to the limited activity of SIRT4, 6, and 7 (Supplemental Figure 2). In the absence of sirtuin enzymes, all p86-Acyl_8,9_ peptides showed less activity than native p86 (Figure 2A). We observed that SIRT1, 2, 3, and 5 restored the activity of modified p86-Acyl_8,9_ peptides in a manner consistent with their reported substrate specificities (Figure 2B). Specifically, incubation of p86-Acetyl_8,9_ with recombinant SIRT1, 2, or 3 restored signal intensity close to that of unmodified p86, confirming deacetylation activity. In contrast, incubation of p86-Acetyl_8,9_ with SIRT5 showed only low luminescent activity, suggesting that SIRT5 is unable to remove the acetyl modification. This is consistent with several recent reports that found SIRT5 to be a weak deacetylase^5,27^, although this is in contrast with an early report^28^. For p86-Crotonyl, SIRT1 and 2 also restored signal, but not as efficiently as for p86-Acetyl_8,9_. SIRT3 and SIRT5 did not restore the signal for p86-Crotonyl. For p86-Succinyl and p86-Glutaryl, incubation with SIRT5 resulted in the restoration of signal, while SIRT1-3 did not, again in agreement with reported substrate specificities^7^.

**Figure 2.**
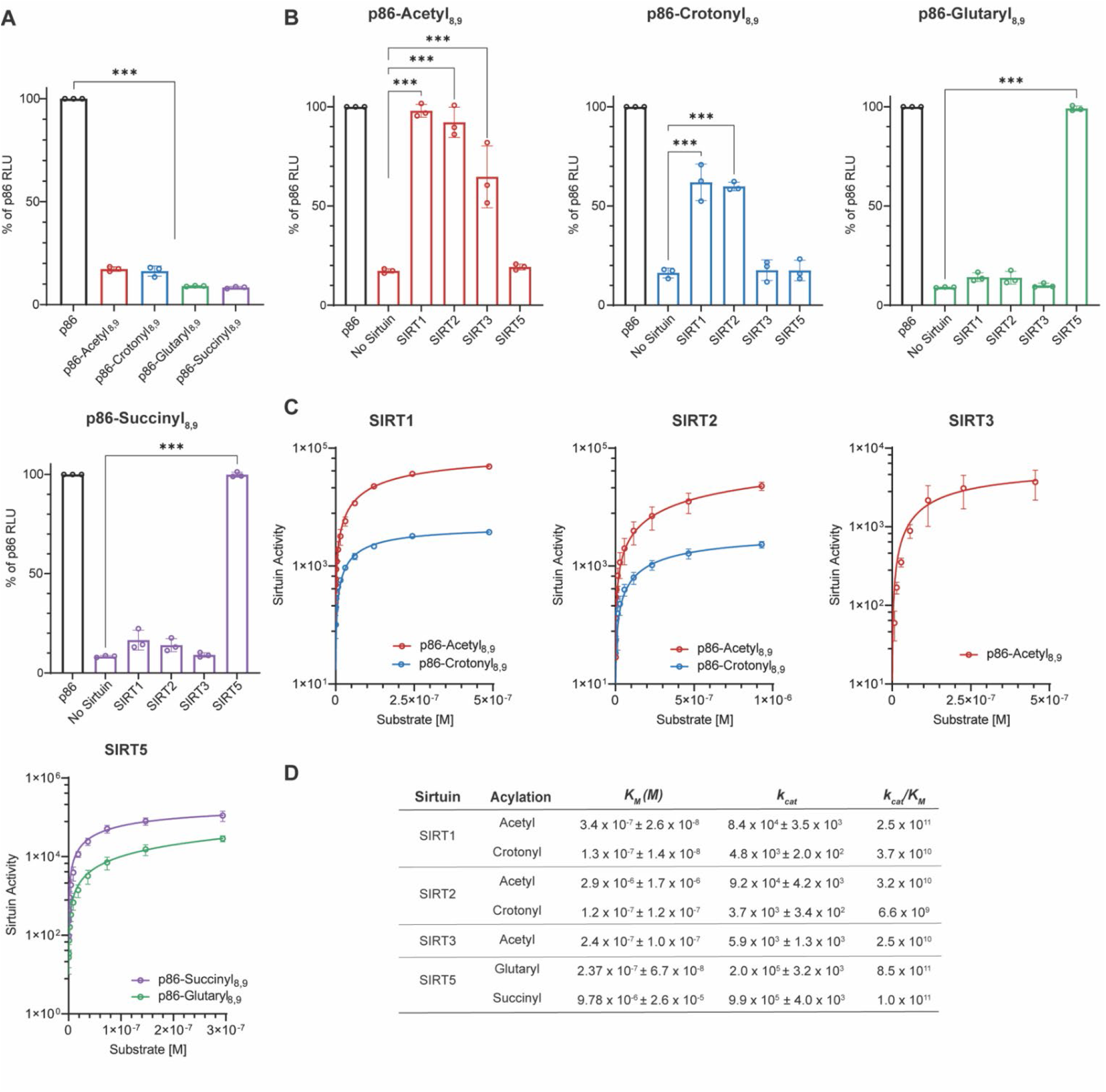
Performance of SIRT*ify* assay in a cell-free system. (**A**) Complementation of unmodified p86 with LgBiT results in luminescence which is significantly decreased for p86-Acyl peptides (-Acetyl, -Crotonyl, -Glutaryl, -Succinyl) modified at K8 and K9. (**B**) The SIRT*ify* assay reveals each sirtuin specificity for different acylations. (**C**) Michaelis-Menten of kinetic parameters of relative kcat and K_M_ for furimazine with sirtuins. (**D**) Table of kinetic parameters determined from curve fits confirms selectivity. Results in **A**–**D** are from three independent experiments. Data show means with standard deviation P values were calculated using one-way ANOVA. \**P* < 0.033, \*\**P* < 0.002, \*\*\**P* < 0.001.

### Sirtuin Deacylation Kinetics of p86-Acylated Peptides

To measure the kinetics of SIRT1, 2, 3, and 5 deacylation, we incubated the sirtuins with acylated p86 peptides (Figure 2C, D). Steady-state deacylation rates were determined by measuring luminescence every five seconds for 10-15 minutes. Comparing deacylation types, SIRT1 and 2 showed *k_cat_* values that were, respectively, ∼24 and ∼15 fold higher for deacetylation compared to decrotonylation. The catalytic efficiencies (*k_cat_/K_M_*) of SIRT1 and 2 were ∼4-6 fold greater for deacetylation than for decrotonylation. For SIRT5, the *k_cat_* values were 5-fold higher for desuccinylation compared to deglutarylation, while, in contrast, the *k_cat_/K_M_* were ∼8-fold lower for desuccinylation compared to deglutarylation due to differences in *K_M_*. Comparing between enzymes, for deacetylation, the *k_cat_* value for SIRT3 was similar to SIRT2, while SIRT1 had a *k_cat_* value ∼4-11 fold higher than SIRT 2 and 3. SIRT1 had ∼5 and 11-fold higher *k_cat_/K_M_* compared to SIRT2 and 3, respectively. For decrotonylation, SIRT1 had a *k_cat_* value ∼2-fold higher and a *k_cat_/K_M_* value ∼4-fold higher than SIRT2. Together, these results indicate that sirtuins display specificity and differential catalytic efficiency that corresponds to deacylation of the modified p86 peptide.

### Using SIRT*ify* to Screen for Sirtuin Inhibitors

Given the involvement of sirtuins in diverse disease states, including cancer, neurodegeneration, cardiovascular disease, and diabetes, there is considerable interest in developing sirtuin inhibitors and sirtuin activating compounds (STACs)^11^. To verify that SIRT*ify* is suitable for the identification of such compounds, we tested a set of well-characterized inhibitors, including EX527 (SIRT1 inhibitor), SirReal2 (SIRT2 inhibitor), 3-TYP (SIRT3 inhibitor), and SIRT5 Inhibitor 1 (S5I1). For initial experiments, we selected inhibitor concentrations higher than all reported IC_50_ values (100 μM for 3-TYP and 1 μM for all other inhibitors) (Figure 3A). Incubation with inhibitor reduced the deacylation-associated signal increase for p86-acetyl_8,9_ in the case of SIRT1-3, for p86-crotonyl_8,9_ in the case of SIRT1-2, and for p86-succinyl_8,9_ and p86-glutaryl_8,9_ in the case of SIRT5, demonstrating enzyme inhibition for all deacylase activities studied, confirming published inhibitor data. However, we found the IC_50_ for SIRT1-3 inhibitors to be higher than previously reported (Figure 3C, Table 1)^29–31^. We postulated that the higher IC_50_ of EX527, SirReal2, and 3-TYP may reflect the higher concentration of NAD^+^ used in our assay (250 μM compared to 170 μM)^29^ and tested additional NAD^+^ concentrations^29–31^. Indeed, using a lower concentration of NAD^+^ decreased the IC_50_ for SIRT1; however, it remained slightly higher than previously published data (Supplemental Figure 3). Comparing IC_50_ values between deacylase types for SIRT5 inhibition, interestingly, we measured an approximately 3-fold lower IC_50_ (0.047 μM) for inhibition of SIRT5’s deglutarylase activity compared to its desuccinylase activity (0.15 μM). S5I is the most potent SIRT5 inhibitor reported thus far, but published data are based exclusively on desuccinylation. The current standard, FLUOR DE LYS® (FDL) assay, does not measure inhibition of SIRT5’s deglutarylase activity; however, comparison between SIRT5’s desuccinylase activity determined from SIRT*ify* and from FDL reported values are within error (Table 1, Supplemental Figure 4)^32^.

**Figure 3.**
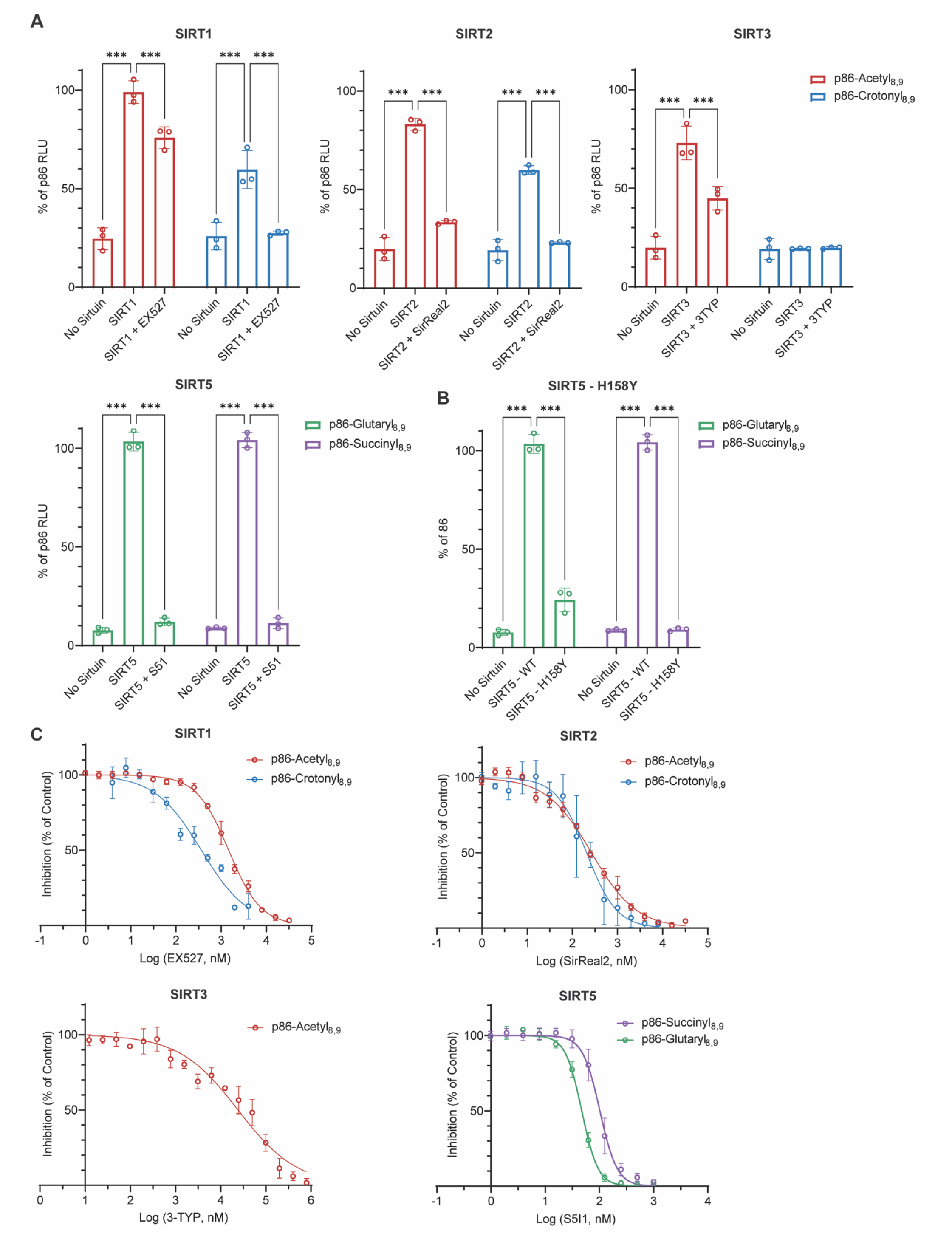
Use of SIRT*ify* to screen sirtuin inhibitors in a cell-free system. (**A**) Specific inhibition of sirtuin activity using small molecule inhibitors. SIRT1; Selisistat (EX527) 1 μM, SIRT2; SirReal2 1uM, SIRT3; 3-TYP 100 μM, SIRT5; SIRT5 Inhibitor 1 (1 μM) (**B**) SIRT5 H158Y mutant activity vs wildtype SIRT5. (**C**) IC_50_ of sirtuin inhibitors against specified substrates. IC_50_ values were estimated by fitting data to nonlinear regression using log(inhibitor) vs. normalized response–variable slope and specified in Table 1. Results in **A**–**D** are from three independent experiments. Data show means with standard deviation. P values were calculated using two-way ANOVA with Tukey’s method of adjustment for multiple comparisons. \**P* < 0.033, \*\**P* < 0.002, \*\*\**P* < 0.001.

**Table 1.**
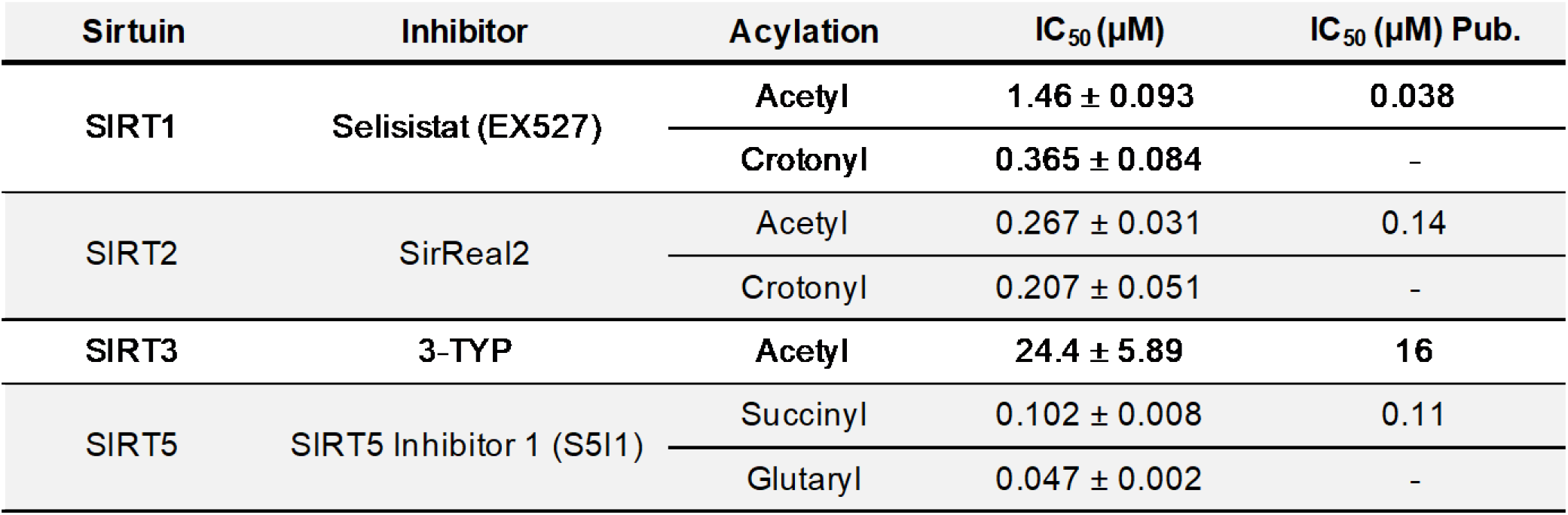
Comparison of IC_50_ values identified from SIRT*ify* to previously published 6913250 findings. References for Selisistat^29^, SirReal2^30^, 3-TYP^31^, SIRT5 Inhibitor 1^32^.

Histidine 158 (H158) is critical for sirtuin catalytic activity^28,33,34^. Replacement of H158 with tyrosine results in a catalytically inactive mutant^28,33,34^. We measured SIRT5-H158Y activity toward p86-succinyl_8,9_ and p86-glutaryl_8,9_. In contrast to wildtype SIRT5, SIRT5-H158Y did not produce any luminescent signal, validating specificity (Figure 3B). In aggregate, these data demonstrate that SIRT*ify* accurately measures inhibitor effects on sirtuin activity and has utility for identifying sirtuin-modulating compounds.

### Measuring Sirtuin Activity in Cells

Currently, available assays for measuring sirtuin activity in cells have significant limitations, including toxicity, availability, complexity of handling due to the need for special safety precautions, and inactivity in many experimental conditions^20–22,35^. To test whether the SIRT*ify* assay can be adapted for use in cells, we stably expressed LgBiT in HepG2 hepatocarcinoma cells and KG1a acute myeloid leukemia cells and focused on SIRT5 (Figure 4A, B). As SIRT5 is primarily located in the mitochondria, we added a mitochondrial targeting sequence (MTS) to LgBiT (LgBiT-MTS) (Figure 4B). As SIRT5 is the main, and possibly the only, mammalian desuccinylase and deglutarylase, we predicted that p86-Succinyl_8,9_ and p86-Glutaryl_8,9_ deacylation activity would be proportional to SIRT5 expression and/or activity. We found that luminescence correlated with p86 concentration. As previously observed, p86-Succinyl_8,9_ had a significantly lower signal as previously observed, however, this change was less dramatic than within the cell-free systems. We first tested whether we can measure changes in SIRT5 activity by NRD167, a cell-permeable S5I1 derivative, or UBSC039, a prospective SIRT5 activator^32,36^. LgBiT-expressing cells were treated with NRD167 or UBSC039 for 2 or 24 hours and then incubated with p86, p86-Succinyl_8,9,_ or p86-Glutaryl_8,9_ in lysis buffer for 2 hours while gently rocking (Figure 4C). Luminescence was measured following the addition of the luminescent substrate furimazine. Consistent with expectations, treatment with NRD167 decreased the luminescent signal with both succinyl and glutaryl-modified peptides in both cell lines, whereas UBSC039 produced a signal increase in only the HepG2 line. Next, we tested whether modifying SIRT5 expression would alter luminescence. We overexpressed (OE) SIRT5 in our LgBiT stably expressing cells and measured luminescence as described above (Figure 4D). SIRT5 OE increased luminescence for both p86-Succinyl_8,9_ and p86-Glutaryl_8,9_ peptides, confirming that SIRT*ify* is sensitive to changes in SIRT5 expression.

**Figure 4.**
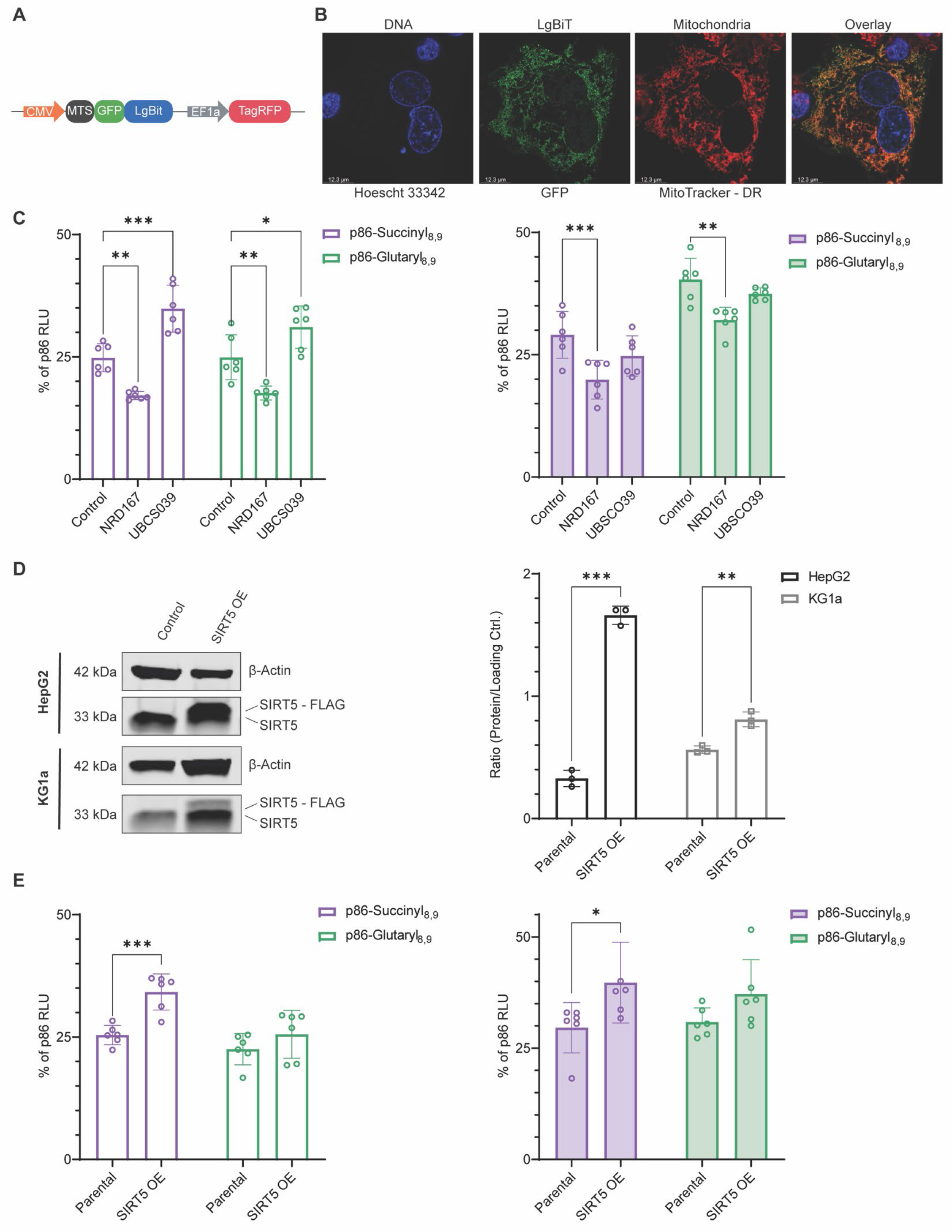
Imaging and quantification of SIRT5 activity in lysed cells. (**A**) Schematic of the pCDH vector transcriptional cassette. **B**) Immunofluorescence imaging of HepG2 cells with stable mitochondrial LgBiT expression (green), nuclei are labeled with Hoechst (blue), and mitochondria with Mitotracker (red). (**C**) HepG2 cells (left) and KG1a cells (right) were treated with the SIRT5 prodrug NRD167 (50 μM) or UBSC0O39 (100 μM) for 2 hours or 24 hours, respectively, before cells were incubated with peptide and lysed. (**D**) Left, analysis of SIRT5 expression in Hep2G cells and KG1a cells. Right, immunoblot quantification. (**E**) HepG2 (left) and KG1a (right) with SIRT5 OE were lysed and treated with peptide for 2 hours. Results in **A**–**D** are from six independent experiments. Data show means with standard deviation. P values were calculated using two-way ANOVA with Tukey’s method of adjustment for multiple comparisons. \**P* < 0.033, \*\**P* < 0.002, \*\*\**P* < 0.001.

We next tested whether SIRT*ify* can be adapted for live-cell measurement of SIRT5 activity. As the utility of a cellular assay is dependent on sufficient cell permeability and distribution, we initially optimized cellular uptake by modifying p86 through C-terminal addition of four arginine residues (p86-R_4_) (Supplemental Figure 5A)^37^. To test whether p86-R_4_ and its derivatives are cell-permeable, we incubated the cell lines with furimazine and graded concentrations of p86-R_4_, p86-Succinyl_8,9_-R_4,_ or p86-glutaryl_8,9-_R4. Luminescence was directly proportional to p86-R4 concentration, while p86-Succinyl_8,9_-R_4_ and p86-Glutaryl_8,9-_R4 had a much lower signal (Supplemental Figure 5B). We first tested whether modification of SIRT5 activity by NRD167 or UBSC039 would modify the deacylase activity of SIRT5 for p86-Succinyl_8,9_-R_4_ and p86-Glutaryl_8,9-_R4 in intact cells. We observed that, consistent with our results on lysed cells, inhibition of SIRT5 resulted in decreased luminescent signal in both cell lines, while UBSCO39 showed an increase only in the HepG2 cell line (Figure 5A). In SIRT5 OE cell lines, we observed increased luminescence in HepG2 and KG1a cells, demonstrating that the utility of the SIRT*ify* assay extends to cell-based screening (Figure 5B).

**Figure 5.**
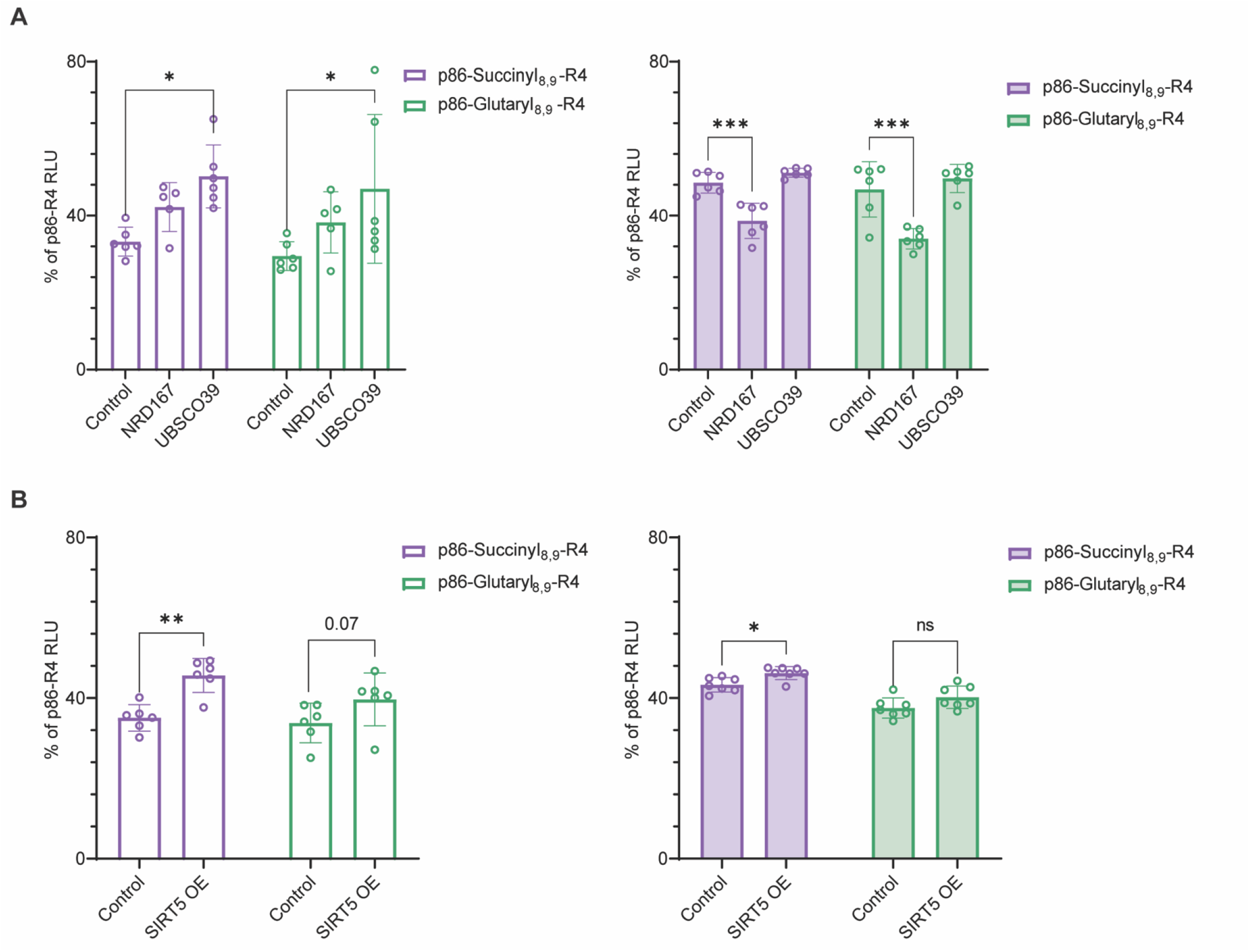
SIRT5 activity in live cells. (**A**) HepG2 cells (left) and KG1a cells (right) were treated with the SIRT5 inhibitor NRD167 (50μM) or UBSCO39 (100μM) for 2 hours or 24 hours, respectively, before incubation with peptide for 2 hours. (**B**) HepG2 (left) and KG1a (right) with SIRT5 OE were treated with peptide for 2 hours. Results in **A**–**B** are from six independent experiments. Data show means with standard deviation. P values were calculated using two-way ANOVA with Tukey’s method of adjustment for multiple comparisons. \**P* < 0.033, \*\**P* < 0.002, \*\*\**P* < 0.001.

### Detecting Sirtuin Activity *In Vivo*

To determine the utility of SIRT*ify* for measuring sirtuin activity in *in vivo*, we injected HepG2 LgBiT-MTS expressing (right) and parental HepG2 (left) cells into opposing flanks of NRG mice (Figure 6A). After the tumor reached ≥5mm size, p86-R4 and furimazine were injected intratumorally, and images were acquired. A strong signal was observed only in flanks injected with HepG2-MTS-LgBiT tumors (Figure 6A). Zebrafish models are a convenient approach to high throughput *in vivo* drug screens. Their major advantage over biochemical and cell line-based screens is that they permit both whole-organism and tissue-specific analysis, potentially accelerating the process of drug development and validation^38^. To test whether SIRT*ify* may be adapted to a zebrafish-based system, we injected purified LgBiT protein with or without p86-R4 peptide into one-cell stage zebrafish embryos, then incubated them with either furimazine or endurazine, a recently reported alternative substrate for live-cell detection that allows for a steady release of furimazine (Supplemental Figure 6). A strong signal was observed in embryos co-injected with p86-R4 and LgBiT but not in embryos injected with LgBiT alone. Next, we tested whether we could measure endogenous *Sirt5* activity in zebrafish. After injecting LgBiT and p86-Succinyl_8,9-_R4, we observed a slight increase in luminescence over the following 90 minutes, suggesting that endogenous desuccinylation activity exists at a low level in early zebrafish embryos and is detected by SIRT*ify* (Figure 6B). Prior studies looking at larval stage zebrafish investigating gain-of-function or loss-of-functions models of *Sirt5* showed that *Sirt5* could substantially contribute to protein succinylation at 7 days post fertilization, suggesting that the modest desuccinylation activity we observed in the first few hours after fertilization might be enhanced at later life stages^39^.

**Figure 6.**
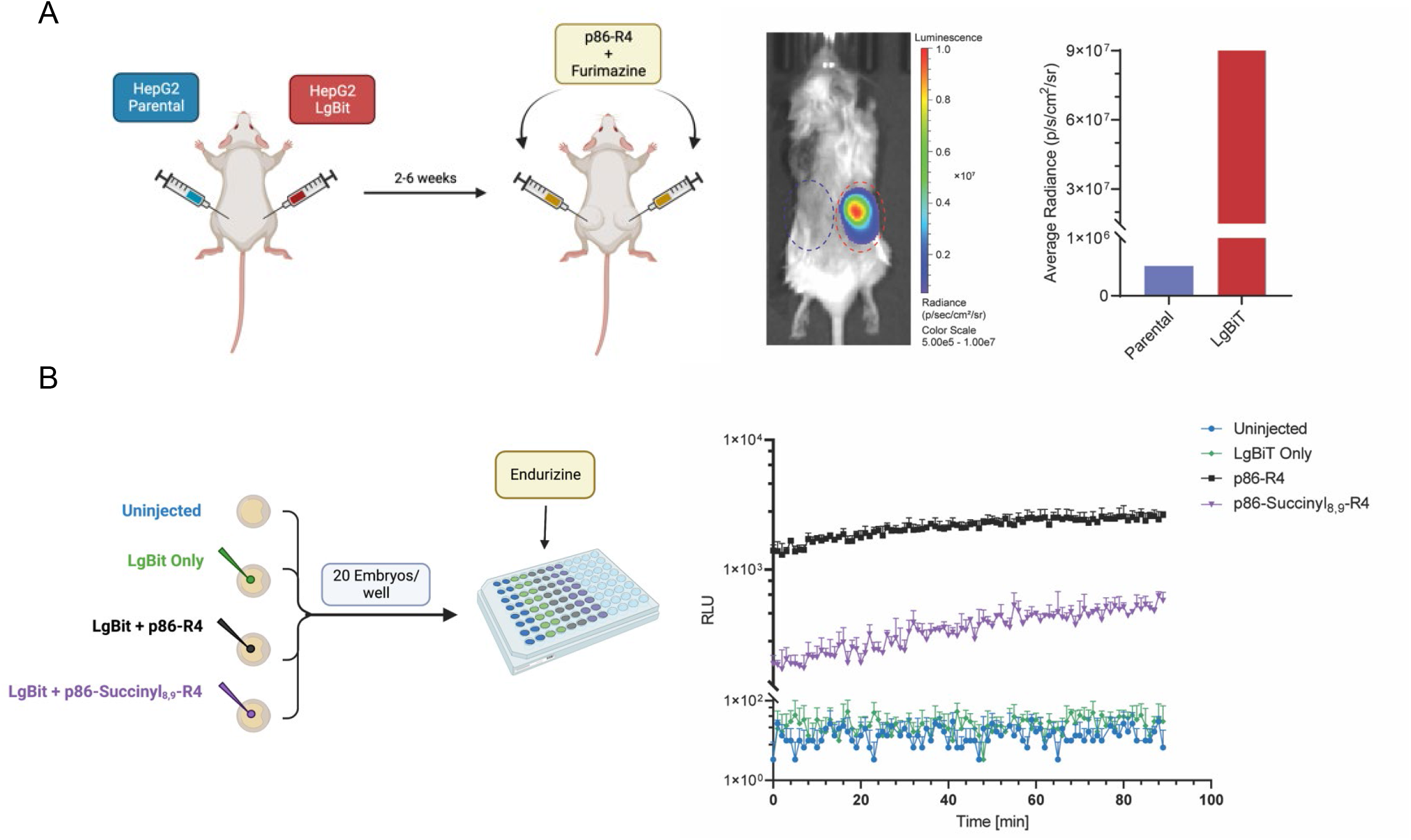
Applicability of SIRT*ify* within *in vivo* systems. (A) Schematic of mouse experiment (left), HepG2 Parental cells were injected into the left flank of an NRG mouse, while HepG2 LgBiT-MTS cells were injected into the right flank. P86 and furimazine were injected into the tumors, and luminescence was measured. Quantification of luminescence (right). (B)Schematic of zebrafish experiment (left), Purified LgBiT with p86 or p86-Succinyl_8,9_-R4 was injected into zebrafish embryos and incubated in the presence of endurazine. Luminescence was measured approximately every 45 seconds for 90 minutes.

## DISCUSSION

Here we describe a novel split-luciferase system that reports sirtuin activity and specificity with high fidelity to benchmarks. The current standard to measure sirtuin activity is the fluorometric system FLUOR DE LYS® (FDL), a two-step process where trypsin is added to cleave the fluorescent AMC conjugate from the deacylated peptide. In cell-free assays and cellular lysates, trypsin may degrade the sirtuin of interest or sirtuin regulatory proteins, a possible explanation for artifacts and lack of specificity^15,16^. We designed SIRT*ify* to overcome this shortcoming, as no second step is required to generate the signal. Additionally, the FDL assay is limited to measuring deacetylation and desuccinylation activity, as there is currently no assay available for other modifications, such as glutaryl and crotonyl. SIRT*ify* not only reproduced previously published results on SIRT1,2,3, and 5 specificities for deacetylation and desuccinylation but also detected decrotonylase and deglutarylase activity, allowing for an easy and quantitative comparison of substrate specificity. It is possible that the same approach could be extended to additional sirtuin activities such as myristoylation or malonylation.

Similar to FDL, assays based on radiolabeled histones, nicotinamide release, fluorescence polarization, or LC-MS are incompatible with live cell measurements^23–25^. Multiple reports have described activity-based chemical probes to measure sirtuin activity in protein mixtures and cell lysates^40,41^. However, additional work is needed to increase the cell permeability, selectivity, and sensitivity of sirtuin activity-based probes (ABPs) before these can be valuable tools to measure sirtuin activity *in vitro*. Here we have demonstrated that SIRT*ify* p86 acyl peptides are cell permeable and can measure changes to SIRT5 activity by either chemical modulators or modifying SIRT5 levels in multiple cell lines. Genetic fluorescent probes have also been described to measure intracellular sirtuin activity. This approach relies on the constitutive expression of an EGFP mutant with a non-canonical acetyl-lysine modification. Where these approaches do not allow for temporally resolved studies (the acetyl-GFP substrate is continuously produced) and are limited to deacetylation activity only, SIRT*ify* can be used for short-or long-term measurements and can detect additional acyl modifications, including succinyl and glutaryl.

p86 does not match any known physiological sirtuin substrate, and there is some evidence that sirtuins may preferentially recognize lysines within a specific sequence context; however, it is challenging to identify a native substrate that is amenable to SIRT*ify*. Reports focused on SIRT2 and SIRT3 failed to demonstrate a clear consensus in the amino acid sequences surrounding the acylated lysine residue^42,43^. Likewise, for SIRT5, no specific sequence has been identified, although certain patterns for acylations are beginning to emerge^44,45^. Differences for preferred substrate acyl groups are caused by binding of the acyl moiety to an active site channel within the respective sirtuin. Although sirtuins share a conserved catalytic core of ∼275 amino acids, differences within the binding cleft configuration distinguish between sirtuins and are central to substrate specificity^46,47^. This selectivity is supported by our SIRT*ify* results, as the sirtuins we tested not only recognized the small p86 peptide but also maintained specificity toward the acyl modification.

A potential confounding factor is sirtuin subcellular localization. SIRT1 is mainly nuclear, SIRT2 is localized to the cytosol, while SIRT3 and SIRT5 are primarily located within the mitochondria. Additionally, subcellular localization of sirtuins can vary during development, in response to stimuli, and in different cell types^48,49^. For example, SIRT5 is primarily located within the mitochondria but has also been observed within the cytoplasm and nucleus. Investigation has focused mainly on the mitochondrial function of SIRT5 and its role in metabolism. However, recent studies have extended the scope to include the role of SIRT5-mediated histone desuccinylation and its impact on disease^50^. We have demonstrated that LgBiT can be targeted to the mitochondria with little impact on cellular health. Other modifications, e.g., to promote nuclear localization, may be helpful to elucidate cell compartment-specific SIRT5 functions.

The recognized importance of sirtuins in cellular homeostasis, aging, and disease has increased interest in developing therapeutic modulators of sirtuin activity. However, the development of such compounds faces major challenges, including limited target specificity and potency^43^. We have shown that SIRT*ify* measurements correlate well with published data^51^, and we demonstrated that SIRT*ify* is highly selective, allows for the determination of steady-state kinetic measurements, and is readily adaptable for high-throughput screening, which could facilitate the discovery and characterization of therapeutic compounds.

Our data indicate the possibility of adapting SIRT*ify* to measure additional post-translation modifications (PTMs), such as ubiquitination and methylation. This may allow quantifying the activity of deubiquitinating enzymes (DUBs) or lysine demethylases (KDM), respectively. DUBS and KDMs are essential regulators of key cellular processes and are involved in autoimmune disorders, cancer, and neurodegeneration^52,53^. In addition to lysines, p86 contains two serines in positions 2 and 11. Common modifications that occur on serine are O-linked glycosylation, methylation, N-acetylation, and phosphorylation, which can regulate catalytic activity ^54^. Alternatively, additional peptides, characterized by Dixon et al., comprise alternative sequences, such as peptide 78, which has two asparagines added on the N-and C-terminus. Asparagine modifications include phosphorylation, hydroxylation, and N-linked glycosylation^26,55,56^. It may be possible to adapt the SIRT*ify* design to measure the activity of these enzymes in a manner similar to that shown here for sirtuins.

While SIRT*ify* provides many advantages over conventional sirtuin activity assays, limitations remain. It is unknown how the activity of recombinantly expressed and purified sirtuins compares to their activity *in situ* or how cellular localization of the sirtuins, p86 peptides, or LgBiT may affect results. We have shown that expressing of LgBiT tagged with a mitochondrial localization signal is tolerated and may be adapted to other cellular compartments. Peptides with cellular localization signals may also improve the SIRT*ify* signal within cells. Secondly, while we demonstrated the ability of SIRT*ify* to measure diacylation activity within *in vivo* systems, endogenous SIRT5 activity appeared to be low in the early zebrafish embryos tested. While sirtuins are expressed in zebrafish, there is limited information about their activity, especially SIRT5, during the early developmental stages. It is possible that SIRT5 activity is low during the first hours post-fertilization when the experiments described here were performed and might be higher at later stages of development. Further analysis of sirtuin activity at different stages of zebrafish development will be needed for optimization of SIRT*ify*. Lastly, the serum stability of p86 and derivatives is unknown, and alternate sequences may be required for extended measurements in animal models. As previous studies have demonstrated that most cell-penetrating peptides display homogeneous distribution, we expect similar tissue distribution of the p86 peptide *in vivo*^37^. Modifications of p86 by the addition of non-natural amino acids or packaging in nanoparticles could be used to improve stability^57,58^.

### Funding and Acknowledgments

This work was supported by the National Institutes of Health (NIH) National Institute of Health (NIH) grant R21 CA256128. Flow cytometry data collection for this publication was supported by the University of Utah Flow Cytometry Core Facility. Sequencing was performed at the DNA Sequencing Core Facility, University of Utah. We acknowledge Cell Imaging Core at the University of Utah for the use of equipment Leica SP8 White light Laser Confocal and thank Dr. Mike Bridges for their assistance in image acquisition. The authors thank Brayden Halverson for the LgBiT-MTS construct and SIRT*ify* name. The authors thank Dr. Rodney Stewart and Amy Kugath for their assistance with the zebrafish experimental design.

## Methods

### Peptide Sequences (Vivitide)

p86: H_2_N-VSGWRLFKKIS-OH

p86-R4: H2N-VSGWRLFKKISRRRR-OH

p86-Acetyl_8,9_: H_2_N-VSGWRLF(K_Ac_)(K_Ac_)IS-OH p86-Acetyl_8_: H_2_N-VSGWRLF(K_Ac_)KIS-OH

p86-Acetyl_8,9_: H_2_N-VSGWRLFK(K_Ac_)IS-OH

p86-Crotonyl_8,9_: H_2_N-VSGWRLF(K_Cro_)(K_Cro_)IS-OH

p86-Glutaryl_8,9_: H_2_N-VSGWRLF(K_Glut_)(K_Glut_)IS-OH

p86-Glutaryl_8,9_-R4: H_2_N-VSGWRLF(K_Glut_)(K_Glut_)ISRRRR-OH p86-Succinyl_8,9_: H_2_N-VSGWRLF(K_Suc_)(K_Suc_)IS-OH

p86-Succinyl_8_: H_2_N-VSGWRLF(K_Suc_)KIS-OH p86-Succinyl_9_: H_2_N-VSGWRLFK(K_Suc_)IS-OH

p86-Succinyl_8,9_-R4: H_2_N-VSGWRLF(K_Suc_)(K_Suc_)ISRRRR-OH

### Cell Culture

All cells were cultured at 37°C in a humidified incubator supplied with 5% CO_2_. HEK293T/17 cells were cultured in Dulbecco’s Minimum Essential Medium (DMEM, ThermoFisher) supplemented with 10% fetal bovine serum (FBS) (Sigma-Aldrich, St. Louis, MO) and 1% penicillin/streptomycin (Invitrogen). HepG2 and KG1a cells were grown in DMEM supplemented with 10% FBS and 100 U/mL penicillin/streptomycin (P/S). Cells were authenticated using the GenePrint 24 kit (Promega) at the DNA Sequencing Core Facility, University of Utah. All cell lines were screened for mycoplasma using the MycoAlert Mycoplasma Detection Kit (Lonza) and were negative.

### Cell-Free Assays

Assays were set up manually in white flat bottom 96-well plates (BRANDplates®; Sigma-Aldrich) at room temperature. All 100 µL reactions were performed in sirtuin buffer, containing 50 mM Tris-HCL, pH 8.0, 137 mM NaCl, 2.7 mM KCl, 1 mM MgCl_2_, 1 mg/mL BSA (Enzo Life Sciences). All reaction mixtures contained 250 nM of sirtuin (SIRT1-3, 5 Reaction Biology, SIRT4,6,7 Sigma-Aldrich), 10 µM furimazine (Promega), and 250 µM NAD^+^ (Enzo Life Sciences), unless otherwise stated, and were prepared in sirtuin buffer. Reaction mixtures were incubated at room temperature for 30 minutes. The reaction was initiated by the addition of 1-250 nM acylated peptide (Vivitide) and a 1:10,000 dilution of LgBiT subunit of the NanoBiT luciferase (Promega) and read using an Envision plate reader (XCite 2105, PerkinElmer).

### Kinetic Measurements

Assays were set up manually in white flat bottom 96-well plates. Peptides and enzymes were serially diluted in sirtuin buffer for 12-18 concentrations. A mixture of furimazine (10 μM) and LgBiT (1:10,000) in sirtuin buffer was added quickly into wells and briefly mixed before luminescence was measured at 5-10 second increments for 10-15 minutes at room temperature. For sirtuin kinetic measurements, acylated peptide was added at 250 nM after the addition of furimazine and LgBiT mixture.

### Lentivirus Production

Plasmids were transfected as a stoichiometric mixture (21 μg) in HEK293T/17 cells using Lipofectamine 2000 and Plus Reagent (Invitrogen) together with psPAX2 (15 μg) (Addgene plasmid #12260; http://n2t.net/addgene:12260; RRID:Addgene_12260) and pVSV-g (10 μg) (Addgene plasmid #132776 ; http://n2t.net/addgene:132776 ; RRID:Addgene_132776)^59^ to generate lentiviral particles. The virus was concentrated with PEG and stored at −80°C.

### NanoLuc Expression in HepG2 and KG1a Cells

One million HepG2 and KG1a cells were plated in standard medium in the presence of polybrene (8 μg/mL) and transfected with a pCDH-CMV-LgBiT-EF1-TagRFP or pCDH-CMV-MTS:LgBiT: GFP-EF1-TagRFP plasmid. Four days after infection, cells were sorted for RFP or RFP/GFP expression and expanded in culture for one week. Luminescence was then measured by adding 10 µM p86-R4 to 50,000 cells in the presence of furimazine and measured using Envision plate reader.

### Immunofluorescence Staining

HepG2-LgBiT expressing cells were grown in regular media in 8-well chamber slides (ThermoScientific). Cells were washed with PBS, then stained with prewarmed (37°C) DMEM without phenol red supplemented with 10% FBS and 1% P/S containing MitoTracker Deep Red probe (ThermoFisher) for 30 minutes at 37°C. Cells were washed with PBS before staining with Hoechst 33342 (ThermoFisher) for 10 minutes at room temperature. Cells were maintained in DMEM solution during imaging.

### Generation of SIRT5 Overexpression Cells

One million HepG2-LgBiT or KG1a-LgBiT cells were plated in standard medium in the presence of polybrene (8 μg/mL) and transfected with pCDH-CMV-SIRT5-FLAG-EF1-CopGFP. Cells were then processed as described above. Cells obtained in this manner were analyzed for SIRT5 expression by immunoblot.

### Inhibition and Activation of Sirtuins

Cell-free assays were set up manually in white flat bottom 96-well plates (BRANDplates®;Sigma-Aldrich) at room temperature. Inhibitors (SIRT1; Selisistat (EX527), SIRT2; SirReal2, SIRT3; 3-TYP, SIRT5; SIRT5 Inhibitor 1) were combined with sirtuin, and NAD^+^, in sirtuin buffer for 30 minutes at room temperature while rocking. Peptide was then added to each well and incubated for 10-30 minutes. Furimazine was added to wells, gently mixed, and read.

For cell-based assays, cells were plated at 50,000 cells per well and allowed to adhere or settle overnight. Cells were washed with warmed PBS and then treated with an inhibitor or activator (see figure for concentration) in complete DMEM media for 24 hours at 37°C. After incubation, cells were washed and then treated with peptide in NP-40 and Halt™ Protease and Phosphatase Inhibitor Cocktail (ThermoFisher) for 10 minutes at room temperature while rocking. Furimazine was added directly before the plate was read. For intact cells, after incubation, cells were washed and then treated with a peptide in DMEM media without phenol red for 2-4 hours at 37°C. Furimazine in DMEM without FBS was added to cells for five minutes before the plate was read. All samples were normalized to p86 peptide.

### Bioluminescence Imaging and Signal Quantification of SIRT5 Activity in Living Cells

The assay was performed manually in black, flat bottom 96-well plates (BRANDplates®;Sigma-Aldrich). Cells were plated at 50,000 cells per well and allowed to adhere or settle overnight. Cells were washed with warmed PBS before the addition of peptide in DMEM (without phenol red) with 10% FBS (ThermoFisher) for 2-4 hours at 37°C. Furimazine was added five minutes before the plate was read.

### Bioluminescence Imaging in *Vivo*

For imaging in mice, unmodified parental HepG2 or HepG2-LgBiT cells were injected into the left/right flank of NOD.Cg-Rag1tm1Mom Il2rgtm1Wjl/SzJ (NRG) mice (Jackson Laboratory, 00779). Once tumors reached ≥5mm, a mixture of p86-R4 or p86-Succinyl_8,9_-R4 and furamizine in a PEG-300 solution (10% glycerol, 10% ethanol, 10% hydroxyproplycyclodextrin, 35% PEG-300 in water) was injected intratumorally. The surface of the skin was wiped after injections. Mice were injected 2-3 minutes apart. Imaging began immediately post-injection. Images were collected every minute under (Low sensitivity settings) Emission filter, open; field of view, 25 cm; f-stop 8; binning, 1 × 1; and exposure time, 1 s. (High sensitivity settings) Emission filter, open; field of view, 25 cm; f-stop 1.2; binning, 2 × 2 and exposure time, 60 s. Exposure times averaged 1 minute for 15-30 minutes. Imaging was performed using IVIS 200 Spectrum, and analysis was performed with Living Image software (Perkin Elmer). All animal studies were approved by the Institutional Animal Care and Use Committee of the University of Utah (Salt Lake City, UT).

### Bioluminescence Detection in Zebrafish

All experiments and husbandry of zebrafish were approved by and conducted in accordance with the Institutional Animal Care and Use Committee (IACUC) at the University of Utah. Adult zebrafish were maintained by the Centralized Zebrafish Animal Research (CZAR) at the University of Utah. Embryos were obtained from crosses of wildtype adult TuAB strain zebrafish (Danio rerio) and injected at the 1-cell stage. For preliminary testing of endurazine (Promega, N2570) and furimazine (Promega, N1120), 5 µL reactions were made by adding 4 µL LgBiT (Promega, N1120), 0.5 µL 100 µM p86-R4 (Vivitide) or 0.5 µL water, and 0.5 µL (10% of reaction volume) 0.5% Phenol Red dye. 100 embryos per treatment group were injected into the yolk with ∼1.5 nL per embryo of the above reactions, and 100 uninjected siblings were set aside for “no luciferase” controls. 20 embryos from each group were plated into a single well of a white, flat bottom 96-well plates (Bandplates®;Sigma-Aldrich) at room temperature in 200 µL 20 mM HEPES buffered E3 media (5 mM NaCl, 0.17 mM KCl, 0.33 mM CaCl2, 0.33 mM MgCl2), pH 7.5, supplemented with 10% FBS with endurazine (1:100) or furimazine (1:50). Following a 30 minute incubation period luminescence was measured using a TECAN M1000 with kinetic cycle of 1-minute interval for a total of 90 minutes. Experiments were performed in triplicate. After establishing endurazine as the appropriate substrate for this application, the above experiment was repeated, only including a reaction containing 10 µM p86-Succinyl_8,9_-R4 to measure SIRT5 activity in early-developing embryos

### Immunoblot Analysis

For immunoblotting, cells were lysed in 1X RIPA lysis buffer (ThermoFisher) containing Halt™ Protease and Phosphatase Inhibitor Cocktail (ThermoFisher). Protein concentration was measured using Pierce BCA Protein Assay Kit (ThermoFisher). Cellular lysates were boiled in Laemmli sample buffer for 10 minutes, separated on Tris-glycine/SDS-PAGE gels (Bio-Rad), followed by transfer to 0.45 μm nitrocellulose membranes (Bio-Rad). Membranes were blocked in 5% non-fat milk in TBST buffer for 1 hour at room temperature, then incubated with primary antibodies for 2 hours at room temperature or overnight at 4°C with gentle rocking. Rabbit monoclonal anti-SIRT5 (D5E11, Cell Signaling) and rabbit monoclonal anti-β-actin (13E5, Cell Signaling) antibodies were used at concentrations of 1:1000 and 1:2000, respectively. Membranes were washed three times for 5 minutes before secondary antibody was added for 1 hour at room temperature with gentle rocking. Secondary antibodies used were IRDye®680LT donkey anti-mouse (926-68022, LI-COR) and IRDye®800CW anti-rabbit (926-32213, LI-COR). Membranes were washed three times before being imaged with an Odyssey Fluorescent Imaging System (LI-COR, Lincoln, NE). ImageJ software was used to analyze the optical density quantification of immunoblots.

### Statistics

All experiments were performed in triplicate, independently. Prism 9 (GraphPad) was used to perform all statistical analyses. Please see the figure legend for the statistical analysis performed. *P* < 0.05 was considered to be statistically significant, except where noted.

## Notes

### Competing Interest Statement

The authors have declared no competing interest.

## REFERENCES CITED

1. Wang, Z.A., and Cole, P.A. (2020). The Chemical Biology of Reversible Lysine Post-translational Modifications. Cell Chem. Biol. 27, 953–969. https://doi.org/10.1016/j.chembiol.2020.07.002.

2. Wagner, G.R., Bhatt, D.P., O’Connell, T.M., Thompson, J.W., Dubois, L.G., Backos, D.S., Yang, H., Mitchell, G.A., Ilkayeva, O.R., Stevens, R.D., et al. (2017). A Class of Reactive Acyl-CoA Species Reveals the Non-enzymatic Origins of Protein Acylation. Cell Metab. 25, 823–837.e8. 10.1016/j.cmet.2017.03.006.

3. Baeza, J., Smallegan, M.J., and Denu, J.M. (2015). Site-Specific Reactivity of Nonenzymatic Lysine Acetylation. ACS Chem. Biol. 10, 122–128. 10.1021/cb500848p.

4. Song, Y., Wang, J., Cheng, Z., Gao, P., Sun, J., Chen, X., Chen, C., Wang, Y., and Wang, Z. (2017). Quantitative global proteome and lysine succinylome analyses provide insights into metabolic regulation and lymph node metastasis in gastric cancer. Sci. Rep. 7, 42053. 10.1038/srep42053.

5. Hirschey, M.D., and Zhao, Y. (2015). Metabolic Regulation by Lysine Malonylation, Succinylation, and Glutarylation. Mol. & Cell. Proteomics 14, 2308 LP – 2315. 10.1074/mcp.R114.046664.

6. Houtkooper, R.H., Pirinen, E., and Auwerx, J. (2012). Sirtuins as regulators of metabolism and healthspan. Nat. Rev. Mol. Cell Biol. 13, 225–238. 10.1038/nrm3293.

7. Kupis, W., Pałyga, J., Tomal, E., and Niewiadomska, E. (2016). The role of sirtuins in cellular homeostasis. J. Physiol. Biochem. 72, 371–380. 10.1007/s13105-016-0492-6.

8. Donmez, G., and Outeiro, T.F. (2013). SIRT1 and SIRT2: emerging targets in neurodegeneration. EMBO Mol. Med. 5, 344–352. https://doi.org/10.1002/emmm.201302451.

9. Bringman-Rodenbarger, L.R., Guo, A.H., Lyssiotis, C.A., and Lombard, D.B. (2018). Emerging Roles for SIRT5 in Metabolism and Cancer. Antioxid. Redox Signal. 28, 677–690. 10.1089/ars.2017.7264.

10. Guarente, L. (2011). Sirtuins, Aging, and Medicine. N. Engl. J. Med. 364, 2235–2244. 10.1056/NEJMra1100831.

11. Bonkowski, M.S., and Sinclair, D.A. (2016). Slowing ageing by design: the rise of NAD+ and sirtuin-activating compounds. Nat. Rev. Mol. Cell Biol. 17, 679–690. 10.1038/nrm.2016.93.

12. Bosch-Presegué, L., and Vaquero, A. (2011). The dual role of sirtuins in cancer. Genes Cancer 2, 648–662. 10.1177/1947601911417862.

13. Longo, V.D., and Kennedy, B.K. (2006). Sirtuins in Aging and Age-Related Disease. Cell 126, 257–268. 10.1016/j.cell.2006.07.002.

14. Roessler, C., Tüting, C., Meleshin, M., Steegborn, C., and Schutkowski, M. (2015). A Novel Continuous Assay for the Deacylase Sirtuin 5 and Other Deacetylases. J. Med. Chem. 58, 7217–7223. 10.1021/acs.jmedchem.5b00293.

15. Madsen, A.S., and Olsen, C.A. (2012). Substrates for Efficient Fluorometric Screening Employing the NAD-Dependent Sirtuin 5 Lysine Deacylase (KDAC) Enzyme. J. Med. Chem. 55, 5582–5590. 10.1021/jm300526r.

16. Marcotte, P.A., Richardson, P.R., Guo, J., Barrett, L.W., Xu, N., Gunasekera, A., and Glaser, K.B. (2004). Fluorescence assay of SIRT protein deacetylases using an acetylated peptide substrate and a secondary trypsin reaction. Anal. Biochem. 332, 90–99. https://doi.org/10.1016/j.ab.2004.05.039.

17. Beher, D., Wu, J., Cumine, S., Kim, K.W., Lu, S.-C., Atangan, L., and Wang, M. (2009). Resveratrol is Not a Direct Activator of SIRT1 Enzyme Activity. Chem. Biol. Drug Des. 74, 619–624. https://doi.org/10.1111/j.1747-0285.2009.00901.x.

18. Xiangyun, Y., Xiaomin, N., Linping, G., Yunhua, X., Ziming, L., Yongfeng, Y., Zhiwei, C., and Shun, L. (2017). Desuccinylation of pyruvate kinase M2 by SIRT5 contributes to antioxidant response and tumor growth. Oncotarget 8, 6984–6993. 10.18632/oncotarget.14346.

19. Kaeberlein, M., McDonagh, T., Heltweg, B., Hixon, J., Westman, E.A., Caldwell, S.D., Napper, A., Curtis, R., DiStefano, P.S., Fields, S., et al. (2005). Substrate-specific Activation of Sirtuins by Resveratrol*. J. Biol. Chem. 280, 17038–17045. https://doi.org/10.1074/jbc.M500655200.

20. Xuan, W., Yao, A., and Schultz, P.G. (2017). Genetically Encoded Fluorescent Probe for Detecting Sirtuins in Living Cells. J. Am. Chem. Soc. 139, 12350–12353. 10.1021/jacs.7b05725.

21. Wang, Y., Chen, Y., Wang, H., Cheng, Y., and Zhao, X. (2015). Specific Turn-On Fluorescent Probe with Aggregation-Induced Emission Characteristics for SIRT1 Modulator Screening and Living-Cell Imaging. Anal. Chem. 87, 5046–5049. 10.1021/acs.analchem.5b01069.

22. Kawaguchi, M., Ikegawa, S., Ieda, N., and Nakagawa, H. (2016). A Fluorescent Probe for Imaging Sirtuin Activity in Living Cells, Based on One-Step Cleavage of the Dabcyl Quencher. ChemBioChem 17, 1961–1967. https://doi.org/10.1002/cbic.201600374.

23. Shao, D., Yao, C., Kim, M.H., Fry, J., Cohen, R.A., Costello, C.E., Matsui, R., Seta, F., McComb, M.E., and Bachschmid, M.M. (2019). Improved mass spectrometry-based activity assay reveals oxidative and metabolic stress as sirtuin-1 regulators. Redox Biol. 22, 101150. https://doi.org/10.1016/j.redox.2019.101150.

24. McDonagh, T., Hixon, J., DiStefano, P.S., Curtis, R., and Napper, A.D. (2005). Microplate filtration assay for nicotinamide release from NAD using a boronic acid resin. Methods 36, 346–350. https://doi.org/10.1016/j.ymeth.2005.03.005.

25. Swyter, S., Schiedel, M., Monaldi, D., Szunyogh, S., Lehotzky, A., Rumpf, T., Ovádi, J., Sippl, W., and Jung, M. (2018). New chemical tools for probing activity and inhibition of the NAD+-dependent lysine deacylase sirtuin 2. Philos. Trans. R. Soc. B Biol. Sci. 373, 20170083. 10.1098/rstb.2017.0083.

26. Dixon, A.S., Schwinn, M.K., Hall, M.P., Zimmerman, K., Otto, P., Lubben, T.H., Butler, B.L., Binkowski, B.F., Machleidt, T., Kirkland, T.A., et al. (2016). NanoLuc Complementation Reporter Optimized for Accurate Measurement of Protein Interactions in Cells. ACS Chem. Biol. 11, 400–408. 10.1021/acschembio.5b00753.

27. Bringman-Rodenbarger, L.R., Guo, A.H., Lyssiotis, C.A., and Lombard, D.B. (2017). Emerging Roles for SIRT5 in Metabolism and Cancer. Antioxid. Redox Signal. 28, 677–690. 10.1089/ars.2017.7264.

28. Nakagawa, T., Lomb, D.J., Haigis, M.C., and Guarente, L. (2009). SIRT5 Deacetylates Carbamoyl Phosphate Synthetase 1 and Regulates the Urea Cycle. Cell 137, 560–570. https://doi.org/10.1016/j.cell.2009.02.026.

29. M., S.J., Rao, P., Lei, X., Thomas, M., Rory, C., S., D.P., and Julie, H.L. (2006). Inhibition of SIRT1 Catalytic Activity Increases p53 Acetylation but Does Not Alter Cell Survival following DNA Damage. Mol. Cell. Biol. 26, 28–38. 10.1128/MCB.26.1.28-38.2006.

30. Rumpf, T., Schiedel, M., Karaman, B., Roessler, C., North, B.J., Lehotzky, A., Oláh, J., Ladwein, K.I., Schmidtkunz, K., Gajer, M., et al. (2015). Selective Sirt2 inhibition by ligand-induced rearrangement of the active site. Nat. Commun. 6, 6263. 10.1038/ncomms7263.

31. Galli, U., Mesenzani, O., Coppo, C., Sorba, G., Canonico, P.L., Tron, G.C., and Genazzani, A.A. (2012). Identification of a sirtuin 3 inhibitor that displays selectivity over sirtuin 1 and 2. Eur. J. Med. Chem. 55, 58–66. https://doi.org/10.1016/j.ejmech.2012.07.001.

32. Rajabi, N., Auth, M., Troelsen, K.R., Pannek, M., Bhatt, D.P., Fontenas, M., Hirschey, M.D., Steegborn, C., Madsen, A.S., and Olsen, C.A. (2017). Mechanism-Based Inhibitors of the Human Sirtuin 5 Deacylase: Structure–Activity Relationship, Biostructural, and Kinetic Insight. Angew. Chemie Int. Ed. 56, 14836–14841. https://doi.org/10.1002/anie.201709050.

33. Schwer, B., North, B.J., Frye, R.A., Ott, M., and Verdin, E. (2002). The human silent information regulator (Sir)2 homologue hSIRT3 is a mitochondrial nicotinamide adenine dinucleotide–dependent deacetylase. J. Cell Biol. 158, 647–657. 10.1083/jcb.200205057.

34. Papanicolaou, K.N., O’Rourke, B., and Foster, D.B. (2014). Metabolism leaves its mark on the powerhouse: recent progress in post-translational modifications of lysine in mitochondria. Front. Physiol. 5.

35. Bonomi, R., Popov, V., Laws, M.T., Gelovani, D., Majhi, A., Shavrin, A., Lu, X., Muzik, O., Turkman, N., Liu, R., et al. (2018). Molecular Imaging of Sirtuin1 Expression–Activity in Rat Brain Using Positron-Emission Tomography–Magnetic-Resonance Imaging with [18F]-2-Fluorobenzoylaminohexanoicanilide. J. Med. Chem. 61, 7116–7130. 10.1021/acs.jmedchem.8b00253.

36. Iachettini, S., Trisciuoglio, D., Rotili, D., Lucidi, A., Salvati, E., Zizza, P., Di Leo, L., Del Bufalo, D., Ciriolo, M.R., Leonetti, C., et al. (2018). Pharmacological activation of SIRT6 triggers lethal autophagy in human cancer cells. Cell Death Dis. 9, 996. 10.1038/s41419-018-1065-0.

37. Sarko, D., Beijer, B., Garcia Boy, R., Nothelfer, E.-M., Leotta, K., Eisenhut, M., Altmann, A., Haberkorn, U., and Mier, W. (2010). The Pharmacokinetics of Cell-Penetrating Peptides. Mol. Pharm. 7, 2224– 2231. 10.1021/mp100223d.

38. MacRae, C.A., and Peterson, R.T. (2015). Zebrafish as tools for drug discovery. Nat. Rev. Drug Discov. 14, 721–731. 10.1038/nrd4627.

39. Gut, P., Matilainen, S., Meyer, J.G., Pällijeff, P., Richard, J., Carroll, C.J., Euro, L., Jackson, C.B., Isohanni, P., Minassian, B.A., et al. (2020). SUCLA2 mutations cause global protein succinylation contributing to the pathomechanism of a hereditary mitochondrial disease. Nat. Commun. 11, 5927. 10.1038/s41467-020-19743-4.

40. Graham, E., Rymarchyk, S., Wood, M., and Cen, Y. (2018). Development of Activity-Based Chemical Probes for Human Sirtuins. ACS Chem. Biol. 13, 782–792. 10.1021/acschembio.7b00754.

41. Goetz, C.J., Sprague, D.J., and Smith, B.C. (2020). Development of activity-based probes for the protein deacylase Sirt1. Bioorg. Chem. 104, 104232. https://doi.org/10.1016/j.bioorg.2020.104232.

42. Bheda, P., Jing, H., Wolberger, C., and Lin, H. (2016). The Substrate Specificity of Sirtuins. Annu. Rev. Biochem. 85, 405–429. 10.1146/annurev-biochem-060815-014537.

43. Dai, H., Sinclair, D.A., Ellis, J.L., and Steegborn, C. (2018). Sirtuin activators and inhibitors: Promises, achievements, and challenges. Pharmacol. Ther. 188, 140–154. https://doi.org/10.1016/j.pharmthera.2018.03.004.

44. Rardin, M.J., He, W., Nishida, Y., Newman, J.C., Carrico, C., Danielson, S.R., Guo, A., Gut, P., Sahu, A.K., Li, B., et al. (2013). SIRT5 Regulates the Mitochondrial Lysine Succinylome and Metabolic Networks. Cell Metab. 18, 920–933. https://doi.org/10.1016/j.cmet.2013.11.013.

45. Nishida, Y., Rardin, M.J., Carrico, C., He, W., Sahu, A.K., Gut, P., Najjar, R., Fitch, M., Hellerstein, M., Gibson, B.W., et al. (2015). SIRT5 Regulates both Cytosolic and Mitochondrial Protein Malonylation with Glycolysis as a Major Target. Mol. Cell 59, 321–332. https://doi.org/10.1016/j.molcel.2015.05.022.

46. Feldman, J.L., Dittenhafer-Reed, K.E., and Denu, J.M. (2012). Sirtuin Catalysis and Regulation *. J. Biol. Chem. 287, 42419–42427. 10.1074/jbc.R112.378877.

47. Rauh, D., Fischer, F., Gertz, M., Lakshminarasimhan, M., Bergbrede, T., Aladini, F., Kambach, C., Becker, C.F.W., Zerweck, J., Schutkowski, M., et al. (2013). An acetylome peptide microarray reveals specificities and deacetylation substrates for all human sirtuin isoforms. Nat. Commun. 4, 2327. 10.1038/ncomms3327.

48. Tanno, M., Sakamoto, J., Miura, T., Shimamoto, K., and Horio, Y. (2007). Nucleocytoplasmic Shuttling of the NAD^+^-dependent Histone Deacetylase SIRT1 *. J. Biol. Chem. 282, 6823–6832. 10.1074/jbc.M609554200.

49. Yanagisawa, S., Baker, J.R., Vuppusetty, C., Koga, T., Colley, T., Fenwick, P., Donnelly, L.E., Barnes, P.J., and Ito, K. (2018). The dynamic shuttling of SIRT1 between cytoplasm and nuclei in bronchial epithelial cells by single and repeated cigarette smoke exposure. PLoS One 13, e0193921.

50. Zorro Shahidian, L., Haas, M., Le Gras, S., Nitsch, S., Mourão, A., Geerlof, A., Margueron, R., Michaelis, J., Daujat, S., and Schneider, R. (2021). Succinylation of H3K122 destabilizes nucleosomes and enhances transcription. EMBO Rep. 22, e51009. https://doi.org/10.15252/embr.202051009.

51. Smith, B.C., Hallows, W.C., and Denu, J.M. (2009). A continuous microplate assay for sirtuins and nicotinamide-producing enzymes. Anal. Biochem. 394, 101–109. https://doi.org/10.1016/j.ab.2009.07.019.

52. Harrigan, J.A., Jacq, X., Martin, N.M., and Jackson, S.P. (2018). Deubiquitylating enzymes and drug discovery: emerging opportunities. Nat. Rev. Drug Discov. 17, 57–78. 10.1038/nrd.2017.152.

53. Shi, Y. (2007). Histone lysine demethylases: emerging roles in development, physiology and disease. Nat. Rev. Genet. 8, 829–833. 10.1038/nrg2218.

54. Shi, Y. (2009). Serine/Threonine Phosphatases: Mechanism through Structure. Cell 139, 468–484. 10.1016/j.cell.2009.10.006.

55. Rodriguez, J., Haydinger, C.D., Peet, D.J., Nguyen, L.K., and von Kriegsheim, A. (2020). Asparagine Hydroxylation is a Reversible Post-translational Modification. Mol. Cell. Proteomics 19, 1777–1789. 10.1074/mcp.RA120.002189.

56. Lee, J.M., Hammarén, H.M., Savitski, M.M., and Baek, S.H. (2023). Control of protein stability by post-translational modifications. Nat. Commun. 14, 201. 10.1038/s41467-023-35795-8.

57. Gentilucci, L., De Marco, R., and Cerisoli, L. (2010). Chemical Modifications Designed to Improve Peptide Stability: Incorporation of Non-Natural Amino Acids, Pseudo-Peptide Bonds, and Cyclization. Curr. Pharm. Des. 16, 3185–3203. http://dx.doi.org/10.2174/138161210793292555.

58. Knauer, N., Pashkina, E., and Apartsin, E. (2019). Topological Aspects of the Design of Nanocarriers for Therapeutic Peptides and Proteins. Pharmaceutics 11, 91. 10.3390/pharmaceutics11020091.

59. Stewart, S.A., Dykxhoorn, D.M., Palliser, D., Mizuno, H., Yu, E.Y., An, D.S., Sabatini, D.M., Chen, I.S.Y., Hahn, W.C., Sharp, P.A., et al. (2003). Lentivirus-delivered stable gene silencing by RNAi in primary cells. RNA 9, 493–501. 10.1261/rna.2192803.

