## Supplemental Material for "Bioluminescence Assay of Lysine Deacylase Sirtuin Activity"

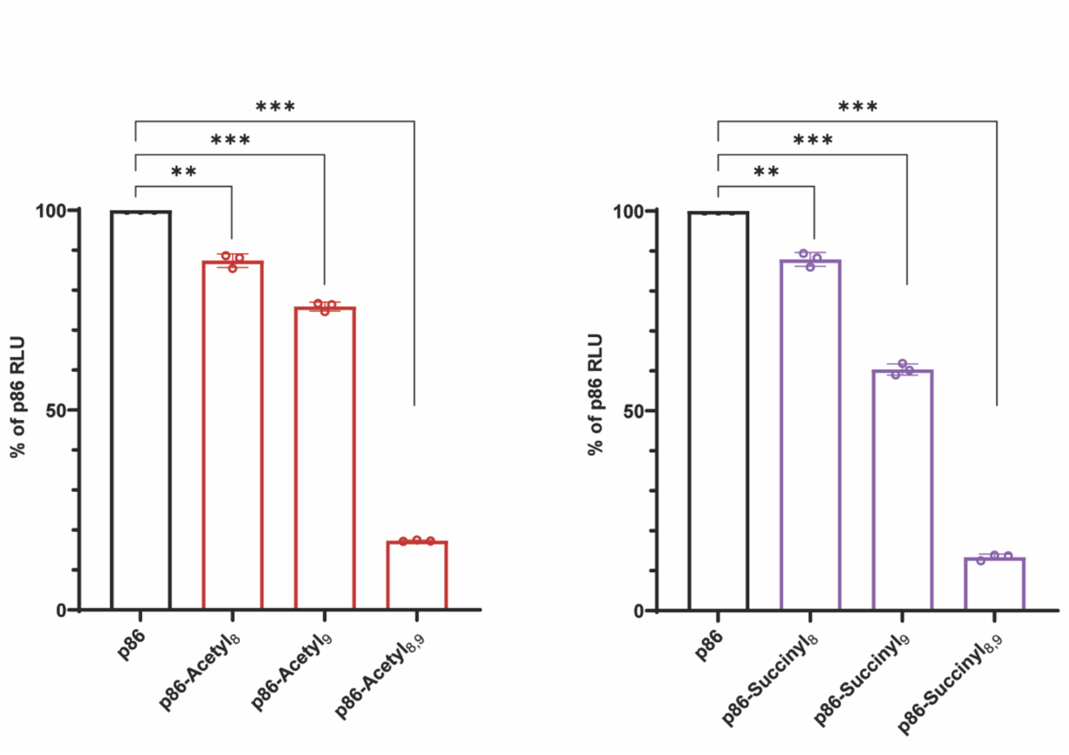


**Supplemental Figure 1.** (A) Comparison of single vs. dual acylation at site K8 and/or K9 of p86 peptide with acetyl (left) or succinyl (right). Total luminescence was measured after the combination of modified p86 peptide with LgBiT. P values were calculated using a paired t-test., $^{*}P < 0.033$, $^{**}P < 0.002$, $^{***}P < 0.001$. Results are from three independent experiments. Data show means with standard deviation.

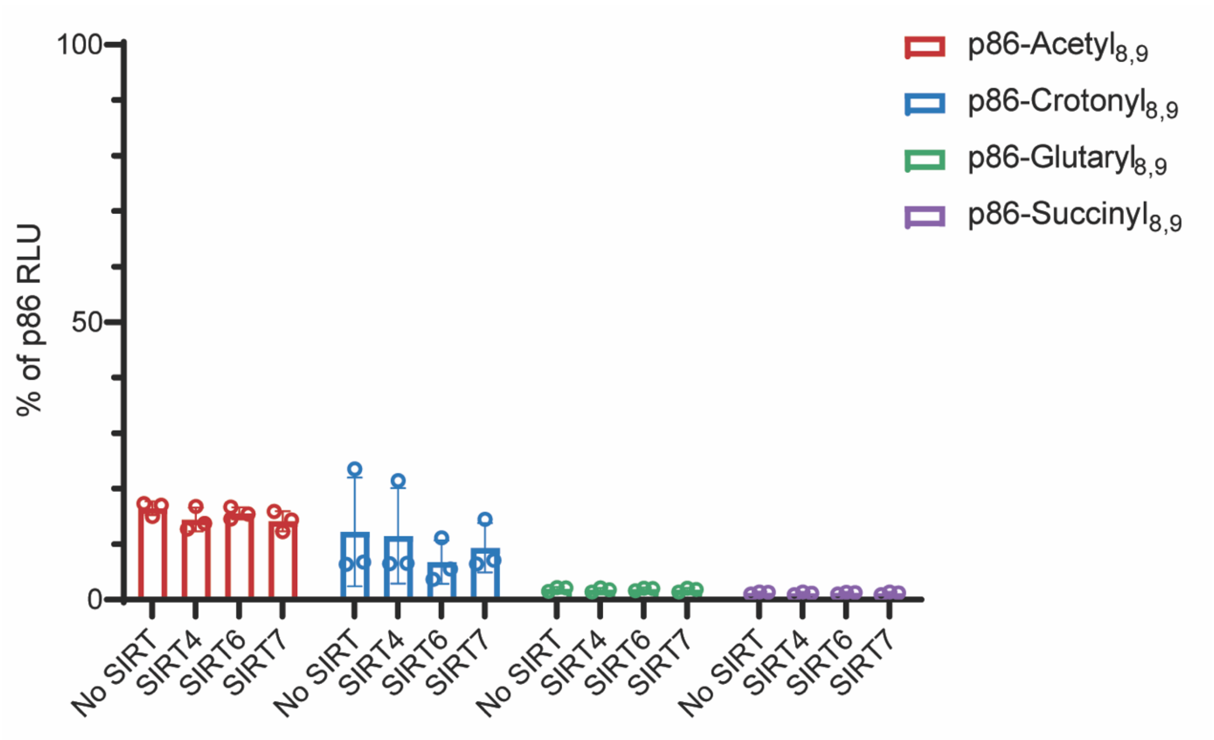


**Supplemental Figure 2.** Deacylation activity of SIRT4, 6, and 7 toward acylated p86 peptides. P values were calculated using two-way ANOVA with Tukey's method of adjustment for multiple comparisons. Results are from independent experiments performed three times. Data show averages with standard deviation.

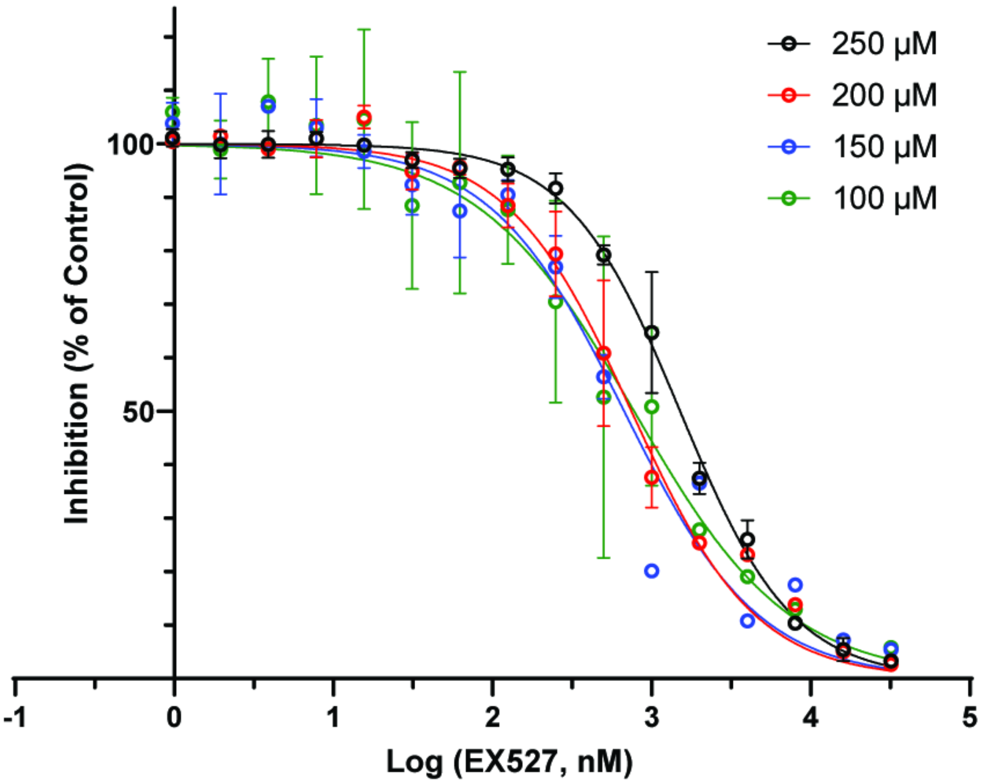


**Supplemental Figure 3.** Response to EX527 in the presence of graded concentrations of $NAD^{+}$. $IC_{50}$ values were estimated by fitting data fitted to nonlinear regression using log(inhibitor) vs. normalized response - variable slope. Results are from three independent experiments. Data show means with standard deviation.

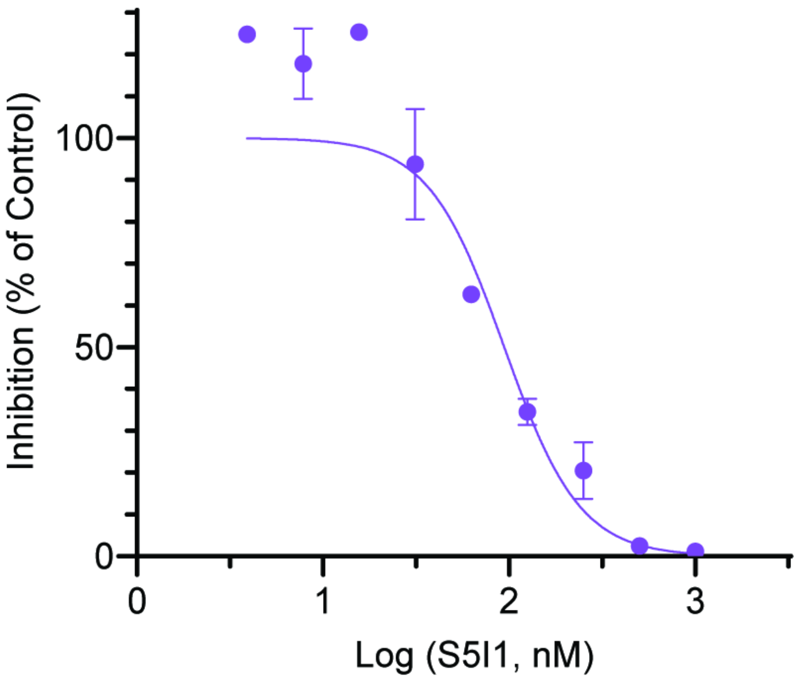


**Supplemental Figure 4.** $IC_{50}$ of SIRT5 inhibitor S5I1 with the commercial Fluor de Lys assay. $IC_{50}$ values were estimated by fitting data fitted to nonlinear regression using log(inhibitor) vs. normalized response - variable slope. $IC_{50}$ = 0.093 nM. Results are from two independent experiments. Data show averages with standard deviation.

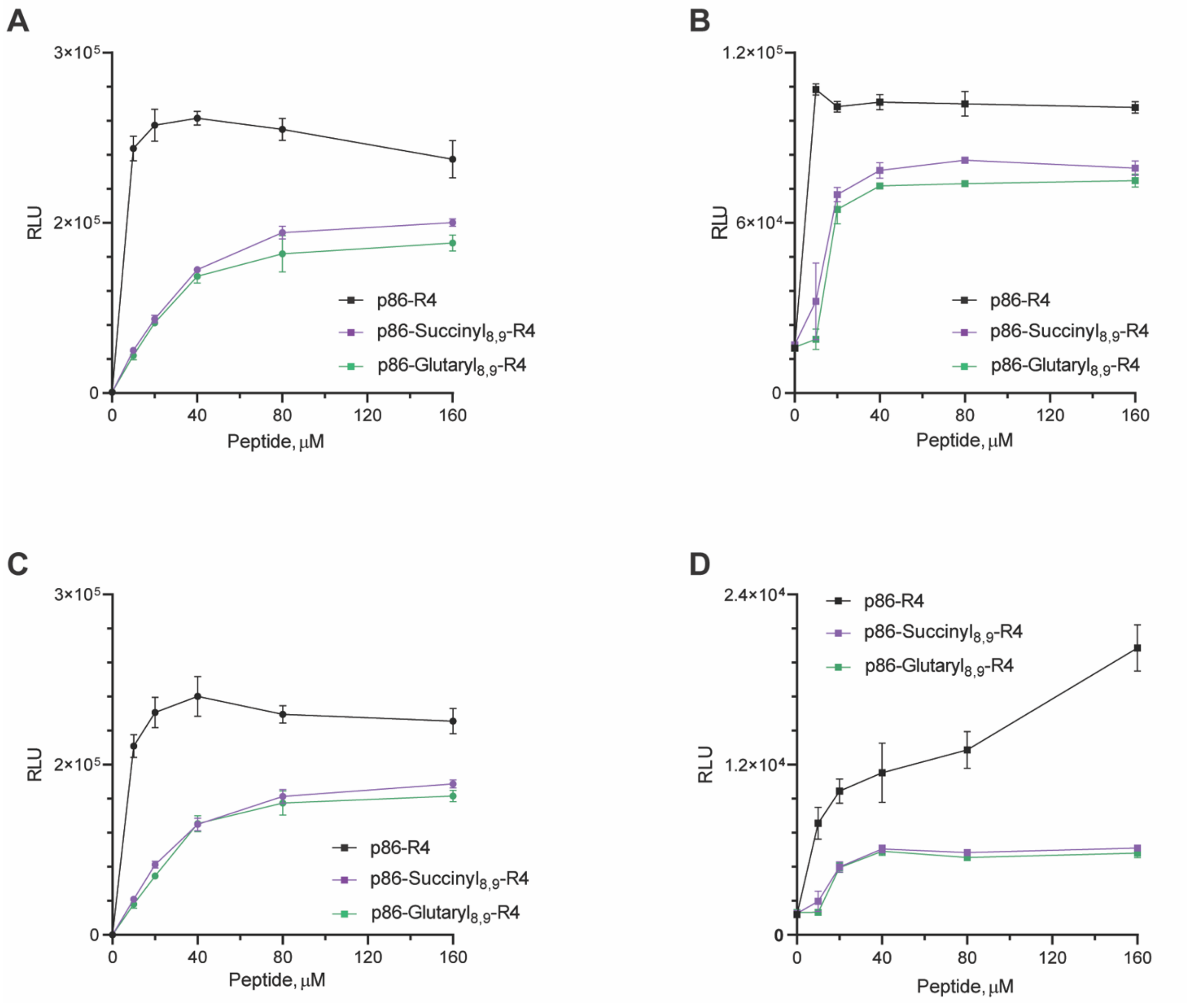


**Supplemental Figure 5.** Peptide titration in HepG2-LgBiT and Kg1a-LgBiT cells. (A-B) Peptide was added to HepG2-LgBiT (A) or KG1a (B) lysate at concentrations from 0-160 µM and incubated for 30 minutes at room temperature. (C-D) Peptide was added to intact HepG2-LgBiT (C) or KG1a (D) cells for 1 hour and incubated at 37°C. Results are from three independent experiments. Data show means with standard deviation.

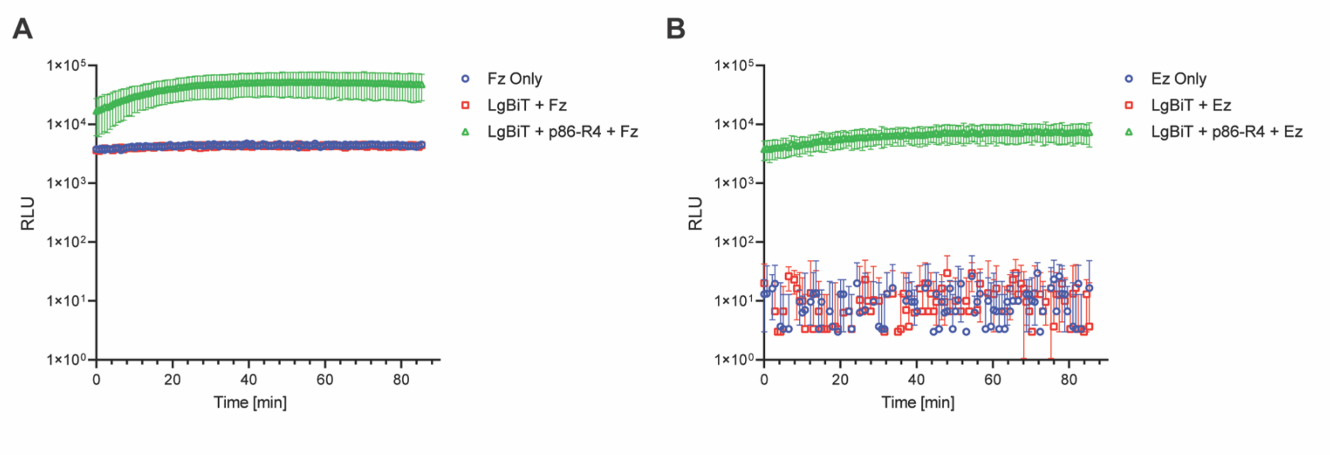


**Supplemental Figure 6.** Substrate comparison in zebrafish with p86-R4 and LgBiT. Twenty zebrafish embryos were injected with a mixture of purified LgBiT with or without p86-R4 and incubated in furimazine (A) or endurazine (B). Embryos were incubated for 20 minutes, then luminescence was measured for 90 minutes. Experiments were performed three times. Data show means with standard deviation.
